# Brain Charts for the Rhesus Macaque Lifespan

**DOI:** 10.1101/2024.08.28.610193

**Authors:** S. Alldritt, J.S.B. Ramirez, R. Vos de Wael, R. Bethlehem, J. Seidlitz, Z. Wang, K. Nenning, N.B. Esper, J. Smallwood, A.R. Franco, K. Byeon, A. Alexander-Bloch, D.G. Amaral, C. Amiez, F. Balezeau, M.G. Baxter, G. Becker, J. Bennett, O. Berkner, E.L.A. Blezer, A.M. Brambrink, T. Brochier, B. Butler, L.J. Campos, E. Canet-Soulas, L. Chalet, A. Chen, J. Cléry, C. Constantinidis, D.J. Cook, S. Dehaene, L. Dorfschmidt, C.M. Drzewiecki, J.W. Erdman, S. Everling, A. Falchier, L. Fleysher, A. Fox, W. Freiwald, M. Froesel, S. Froudist-Walsh, J. Fudge, T. Funck, M. Gacoin, D.J. Gale, J. Gallivan, C.M. Garin, T.D. Griffiths, C. Guedj, F. Hadj-Bouziane, S.B. Hamed, N. Harel, R. Hartig, B. Hiba, B.R. Howell, B. Jarraya, B. Jung, N. Kalin, J. Karpf, S. Kastner, C. Klink, Z.A. Kovacs-Balint, C. Kroenke, M.J. Kuchan, S.C. Kwok, K.N. Lala, D.A. Leopold, G. Li, P. Lindenfors, G. Linn, R.B. Mars, K. Masiello, R.S. Menon, A. Messinger, M. Meunier, K. Mok, J.H. Morrison, J. Nacef, J. Nagy, V. Neudecker, M. Neuringer, M.P. Noonan, M. Ortiz-Rios, J.F. Perez-Zoghbi, C.I. Petkov, M. Pinsk, C. Poirier, E. Procyk, R. Rajimehr, S.M. Reader, D.A. Rudko, M.F.S. Rushworth, B.E. Russ, J. Sallet, M.M. Sanchez, M.C. Schmid, C.M. Schwiedrzik, J.A. Scott, J. Sein, K.K. Sharma, A. Shmuel, M. Styner, E.L. Sullivan, A. Thiele, O.S. Todorov, D. Tsao, A. Tusche, R. Vlasova, Z. Wang, L. Wang, J. Wang, A.R. Weiss, C.R.E. Wilson, E. Yacoub, W. Zarco, Y. Zhou, J. Zhu, D. Margulies, D. Fair, C. Schroeder, M. Milham, T. Xu

## Abstract

Recent efforts to chart human brain growth across the lifespan using large-scale MRI data have provided reference standards for human brain development. However, similar models for nonhuman primate (NHP) growth are lacking. The rhesus macaque, a widely used NHP in translational neuroscience due to its similarities in brain anatomy, phylogenetics, cognitive, and social behaviors to humans, serves as an ideal NHP model. This study aimed to create normative growth charts for brain structure across the macaque lifespan, enhancing our understanding of neurodevelopment and aging, and facilitating cross-species translational research. Leveraging data from the PRIMatE Data Exchange (PRIME-DE) and other sources, we aggregated 1,522 MRI scans from 1,024 rhesus macaques. We mapped non-linear developmental trajectories for global and regional brain structural changes in volume, cortical thickness, and surface area over the lifespan. Our findings provided normative charts with centile scores for macaque brain structures and revealed key developmental milestones from prenatal stages to aging, highlighting both species-specific and comparable brain maturation patterns between macaques and humans. The charts offer a valuable resource for future NHP studies, particularly those with small sample sizes. Furthermore, the interactive open resource (https://interspeciesmap.childmind.org) supports cross-species comparisons to advance translational neuroscience research.

## Main

The rhesus macaque is among the most extensively utilized nonhuman models in translational neuroscience, owing to its resemblance in brain anatomy, phylogenetics, cognitive, and social behaviors to humans. Beyond the study of similarities and differences across species, which have led to key insights into the evolution and organization of human brain function, researchers are increasingly turning to nonhuman primate (NHP) models to gain mechanistic insights into brain development. Toward this goal, there are growing ambitions to establish population models of normative neurodevelopment in rhesus macaques across the lifespan, as this would provide insights into the etiology of pathological human brain development, as well as contribute to our understanding of brain vulnerability, plasticity, and aging^1–3^. The study of neurodevelopment in NHPs offers distinct advantages over similar longitudinal human studies^3^ in two key regards - a shorter lifespan (macaques typically live 25-30 years in captivity), which allows for more rapid generation of life-spanning data compared to humans, and the feasibility for conducting controlled experimental manipulations, particularly in pediatric research to target developmental stages for potential intervention.

The human neuroimaging literature has recently provided a normative trajectory model for accelerating and scaling the generation of growth charts for brain structure over the lifespan^4^. Leveraging advancements in modern imaging techniques and, most importantly, the increasing availability of openly shared datasets that collectively cover the lifespan, this recent effort has quantified age-related changes in the human brain and generated growth charts using an aggregate dataset of over 120,000 scans^4^. However, we still lack comparative benchmark studies in NHPs, with a significant barrier being the establishment of normative growth charts that include a robust cohort of animals across their entire lifespan^5–7^. Despite this lag, there has been important progress in the field of NHP research. Data sharing in NHPs has emerged as a means of overcoming traditionally small sample sizes in this domain, spurring the development of tools that are robust to the many nuances of preprocessing for NHP imaging data, as well as variations in study procedures (e.g., anesthesia, scanner hardware), and heterogeneous magnetic resonance imaging (MRI) protocols used for data acquisition^8^. Additionally, an increasing number of studies have begun to sample developmental and aging populations, creating opportunities for collectively mapping brain changes across the lifespan in a robust manner. In particular, a recent open science initiative - PRIMatE Data Exchange (PRIME-DE)^9–12^ - has collectively aggregated data from over 1000 NHPs, providing the opportunity to bridge this gap and establish comparative NHP brain growth charts across the lifespan.

In this study, we aimed to build the corresponding normative growth charts for brain structure over the macaque lifespan that mirror those already generated for humans^4^. To achieve the sample necessary for this seemingly lofty task of an NHP model, we collectively aggregated over 1,500 MRI scans across more than 1,000 rhesus macaques shared through PRIME-DE and other research studies (see Supplementary Information S1.1-9). Similar to the human brain charts^4^, we used the generalized additive model for location, scale, and shape (GAMLSS) framework to build brain developmental trajectories of brain volume, cortical thickness, and surface area. Furthermore, we compared the development of global and regional brain structures between macaques and humans, identifying corresponding developmental milestones from the prenatal stage to aging. Finally, we delineated species-specific brain maturational changes and critical development stages for brain networks and relevant cognitive functions. We aim to provide: (i) an open resource of sex-stratified growth charts for macaque brain structures that can be utilized as normative trajectories in future NHP studies, particularly for benchmarking studies with small NHP samples; and (ii) age-related, species-comparable neurodevelopmental trajectories to support future translational efforts between human and macaques.

### Normative models of macaque brain growth

Consistent with the approach previously employed in humans, we built global and regional brain growth charts for macaques across the lifespan using GAMLSS^14^. GAMLSS is a widely used distributional regression model, where parameters of location (i.e. mean), variation, skewness, and kurtosis or tails can be modeled as a function of age and sex in developmental studies to establish the inferred non-linear normative trajectories^15,16^. Leveraging large-scale datasets across many macaque studies, sex-stratified and age-related changes were modeled by optimizing fractional polynomial terms in GAMLSS (see Methods for details of model generation and optimization). To account for site-specific batch effects across studies, ‘site’ was included in GAMLSS as a random effect parameter. Additionally, the ComBat batch correction approach was applied to harmonize data across differences in preprocessing pipelines^17^. We present details of data acquisitions (Supplementary Information 1), preprocessing (Supplementary Information 2.1), quality control (Supplementary Information 2.2), batch effect correction (Supplementary Information 2.3), model evaluation (Supplementary Information 3), and robustness analyses (Supplementary Information 3). Global and regional structural MRI-derived phenotypes, including tissue volume, cortical thickness, and cortical area were modeled in GAMLSS.

Similar to human development, the trajectories for total gray matter volume (GMV), white matter volume (WMV), and subcortical gray matter volume (sGMV) of macaque brains exhibit significant increases early in life (Fig 1c-d). Specifically, GMV peaks at almost 9 months (0.74 years) of life with a 95% bootstrapped confidence interval (CI) of 0.65 - 0.86 years for both males and females, followed by a slight decrease and a gradual plateau, indicating an early surge in GM development during the infant stage^18^. WMV also demonstrates a rapid increase during the prenatal period, though this was followed by an initial plateau at approximately 3-4 months before finally showing a steady increase throughout the lifespan. In contrast to GMV and WMV, total sGMV exhibited a more gradual increase from birth, peaking during macaque adolescence at 4.39 years in males and females (95% bootstrapped CI: 3.70 - 5.75 years), followed by a steady decline. Conversely, the lateral ventricle shows a distinct pattern, with an initial decrease leading up to birth, followed by steady, linear-like growth during adulthood and an acceleration at the late elder stage. Overall, these findings were consistent across research sites (Fig 1e) and aligned with previous smaller-scale studies^18,19^. Of note, macaques differed more substantively in peak growth rates observed across tissue compartments in humans (Fig 1f), with macaques reaching peak growth rates prenatally, whereas humans reached peak growth rates after birth (i.e. infancy for GMV and sGMV; toddler for WMV).

**Fig 1.**
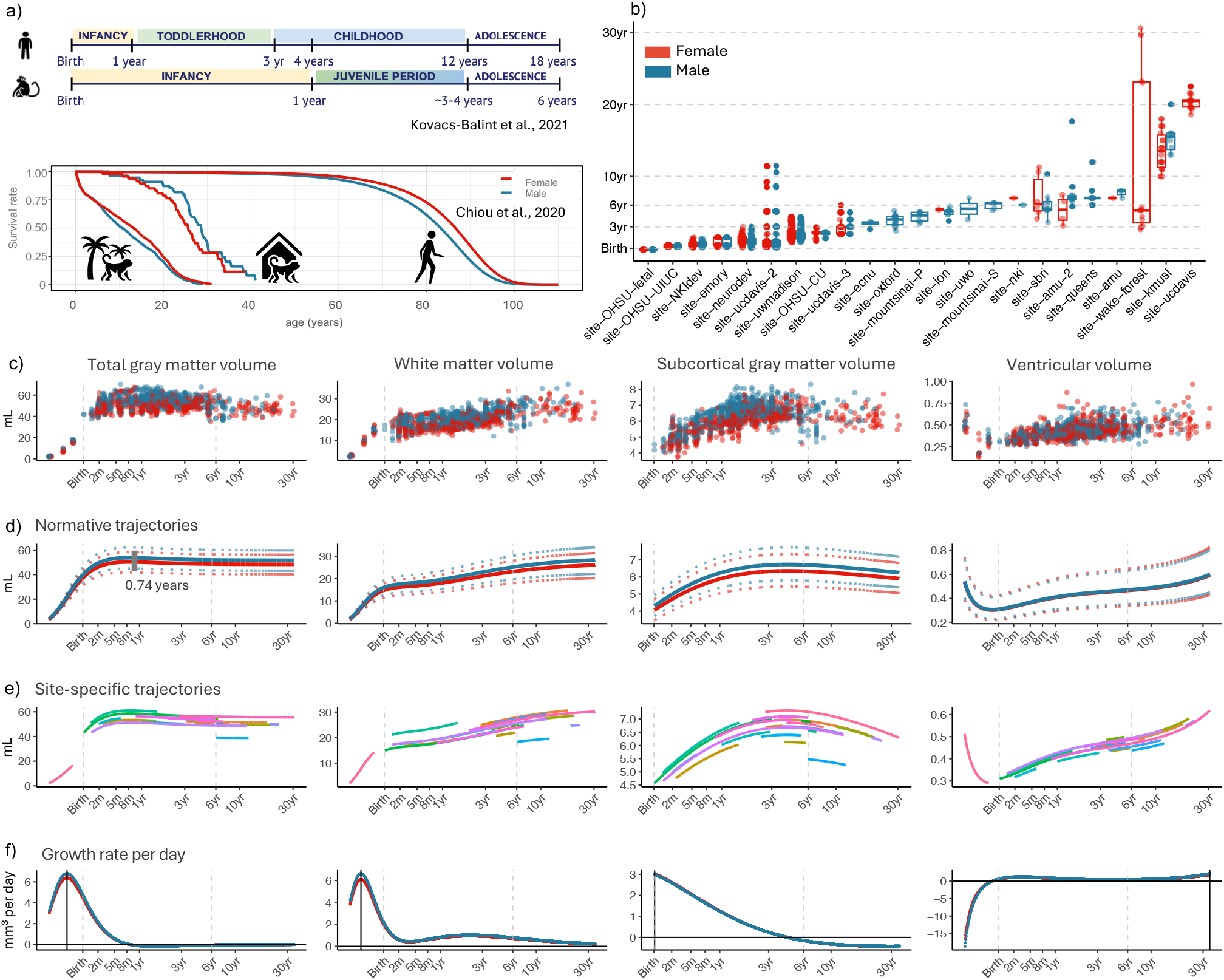
Macaque brain charts. **a**) Comparison of lifespan and age milestones between macaques and human. **b**) Demographic box plot of rhesus macaque data used in the analysis. Data includes 1,024 macaques with 1,522 MRI scans collected from 23 unique sites. **c**) Raw total tissue volume (in *mL*) for cortical gray matter, white matter, subcortical gray matter, and the lateral ventricles. Age (x-axis) is shown on a log scale, with points colored by sex. **d**) Normative trajectories of the four tissue volumes fit with GAMLSS, incorporating sex stratification and site-specific random effects, represented as median centile (the 50th centile) fitted curves of the population. Dotted lines represent 2.5% and 97.5% centile distributions, with both curves colored by sex. **e**) Site-specific curves for each tissue type. Trajectories show ranges of each contributing dataset with the site-specific random effect removed. **f**) Volumetric rate of growth (in *mm*^*3*^ per day) for each of the four global tissue types, colored by sex. Solid vertical lines denote the age at which the growth rate peaked.

Beyond the total volumes of the four tissue types, we also modeled the trajectories for total cerebral volume (TCV), total cortical area (TCA), and mean cortical thickness (Fig 2a-c). Both total cerebral volume and cortical area showed substantial increases after birth, followed by a more gradual, steady increase until an initial plateau phase around 8 months to 1 year, with minimal changes thereafter. Conversely, mean cortical thickness peaks earlier than volume and area metrics, reaching maturity around 3-4 months (0.28 years, CI: 0.23 - 0.30 years), followed by a gradual decline over the lifespan. Similar trajectories were observed in humans, with cortical thickness peaking earlier during infancy, while TCV and TCA reach their maximum later during adolescence. This finding suggests that although cortical thickness plateaus at 3-4 months and begins thinning thereafter, gray matter volume continues to grow for up to approximately 1 year in macaques. Since cortical gray matter volume is a compound measure of its surface area and thickness, the increases in total volume during this period can be attributed to greater area expansion for both humans and macaques.

**Fig 2.**
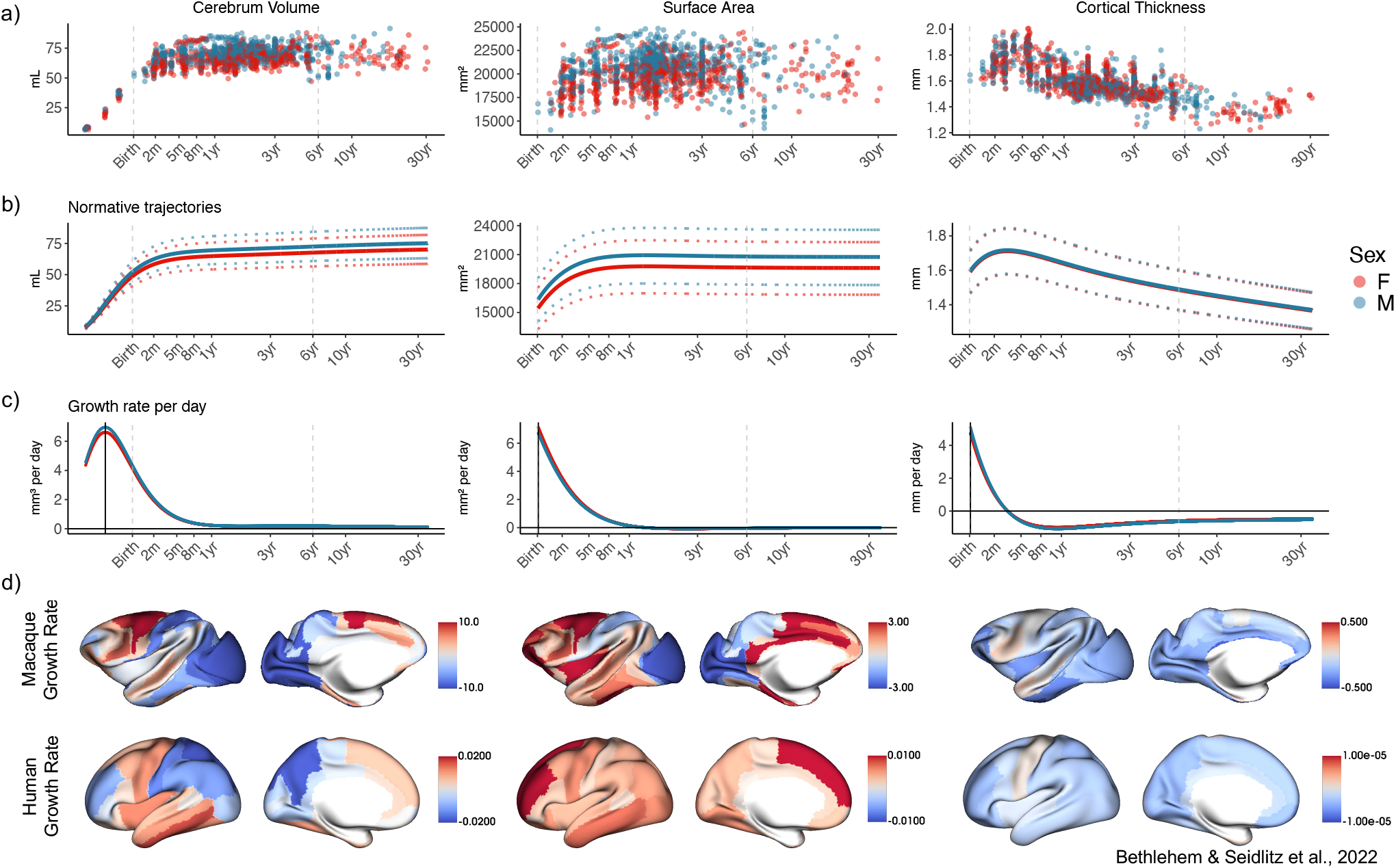
Global and regional cortical morphometric phenotypes and growth rates. **a**) Scatterplots for total cerebrum volume, total cortical surface area, and mean cortical thickness, colored by sex. Volume is measured in *mL*, surface area in *mm*^2^ and cortical thickness in *mm*. All plots are shown on a log-scaled x axis. **b**) Normative trajectories with associated 2.5% and 97.5% centile distributions for each metric, stratified by sex. **c)** Growth rates per day of the respective phenotype, stratified by sex. Solid vertical lines again denote peak growth. **d)** Surface maps of regional growth rate for macaque and human at their respective total gray matter volume peak age (0.74 years for macaque, 5.90 years for human). Sex-stratified models were generated for bilateral averages of each region, and growth rates for male are shown on the left hemisphere for visualization (female maps in Supplementary Information 4.3). Macaque regions are defined by the Markov parcellation and human regions are defined by the Desikan-Killiany parcellation^13^. Human growth rate data was obtained from the human brain charts project^4^.

We characterized regional growth using the Markov architectonic parcellation^20^ (Supplementary Figures 4.1.1). Figure 2d depicts growth velocities of regional gray matter volume, surface area, and cortical thickness at the age of peak total gray matter volume in macaques (i.e. 0.74 years). Additionally, we present the human growth rate at the age of peak GMV estimated in the previous human chart study (i.e. 5.90 years)^4^. When compared in this manner, humans and macaques exhibited similar patterns of regional variation in growth rate for all three morphometric measurements. Volumetrically, in both humans and macaques, the precentral motor cortex continues to grow while a decline is seen in the postcentral sensory cortex, parietal, and occipital lobes. In the lateral frontal regions, both species have reached their peaks and have already started to decline. In contrast, the medial frontal regions take longer to fully mature and continue to grow.

A notable discrepancy between humans and macaques is observed in the insular and ventral temporal cortex, with humans exhibiting strong volumetric growth at the respective peak age of GMV, while the macaque has already begun to decline. For surface area, the entire cortex undergoes cortical expansion in humans at the peak age of GMV. A similar area expansion is observed across the entire macaque cortex, except in the primary visual area, which experienced a decline. Both humans and macaques showed high cortical expansion in the middle cingulate cortex, while macaques showed greater expansion in the precentral motor cortex compared to the postcentral sensory cortex. There is a notable difference in surface area expansion in the prefrontal cortex between species. Humans show rapid increases in total area in this region even after volume and thickness have begun to decline, suggesting a substantial expansion of the frontal surface during this period. In comparison, while the macaque prefrontal cortex is still expanding, it is not growing in area as disproportionately relative to the rest of the brain. Overall, regions vary in volume and area growth in both species, whereas cortical thickness peaks before GMV and begins to thin in most cortical regions.

### Developmental milestones across species

To facilitate cross-species comparison of developmental milestones for brain structure, we generated tissue-specific proportional trajectories in macaques and compared them with those established in the human study (Fig 3). Lifespan milestones in macaques, including gestation, infancy, juvenile, adolescence, adulthood, and elderhood were defined primarily based on key developmental stages and physiological, and behavioral changes. In this context, human age scales to be approximately 3-4 times than that of macaques^19^. In macaques, the growth rates of all gray matter and white matter measurements peaked prenatally - with macaques being born with 55 - 75% of their maximum volumes. In contrast, humans are born with only 25 - 30% of their maximum volume at birth, with the highest growth rates for gray matter occurring during infancy and white matter during toddlerhood. This observation aligns with previous studies, suggesting that humans undergo a unique, extended period of gestation that extends beyond birth^29,30^. The surface maps in Figure 3 show the average regional proportion of gray matter volume at different developmental stages for humans and macaques, respectively. Overall, both species reached their regional maximal volume along the posterior-to-anterior axis, with posterior regions maturing first, followed by anterior (Figure 3 and Supplementary Information 4). During infancy, frontal and temporal lobes in humans only reach 50-60% of their maximum, whereas in macaques at 4 months of age, these regions reach over 90% of their maximum. In both species, gray matter volume starts to decline in the frontal and parietal lobes before decreases are detected in total GMV. Notably, the precentral motor cortex in macaques remains almost the same size throughout the entire macaque lifespan, while age-related reductions are noted in humans, they are notably more moderate than in other regions.

**Fig. 3.**
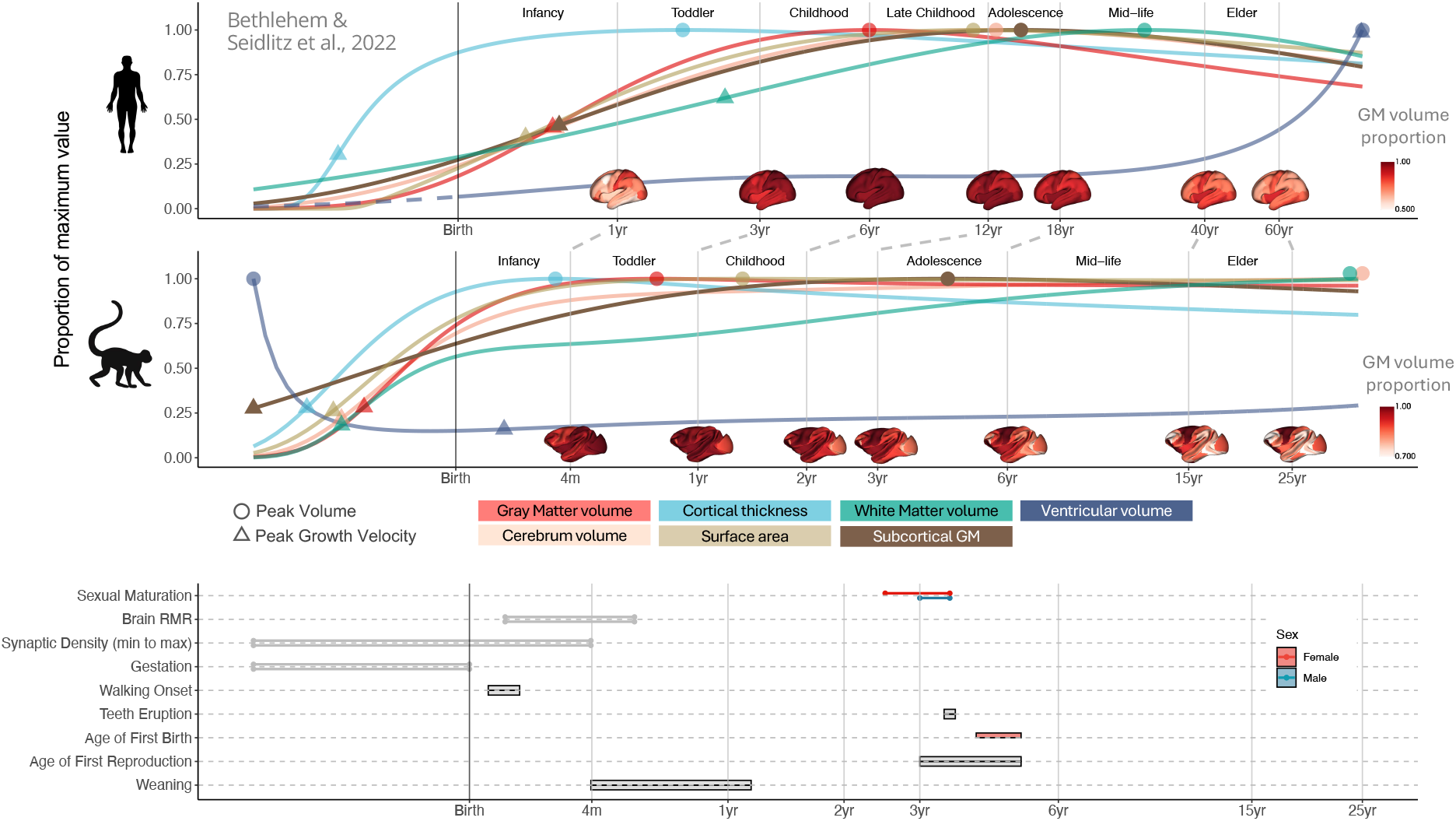
Cross-species comparison of neurodevelopmental milestones. Normative trajectories of each global MRI phenotype for both human (top) and macaque (middle) as a function of age (log-scaled). Regional gray matter volume are shown at highlighted developmental milestones^3,21–28^. Both trajectories and regional gray matter volume are shown as a proportion of their maximum. Circles on the trajectories indicate the peak value of each phenotype (i.e. the maxima of the median trajectories), and triangles represent the peak growth velocity. Developmental milestones are approximately matched for each species for visualization purposes. Bottom panel depicts commonly used developmental milestones in macaque stratified by sex. Ranges depict time of expected onset (bars: walking onset, teeth eruption, age of first birth, first reproduction, weaning) or completion of specific behavioral or physiological maturity (lines: sexual maturation, brain resting metabolic rate (RMR), synaptic density, gestation)^3,21– 28^.

Comparing the trajectories of cortical gray matter development between species, while the order of cortical maturation is consistent (i.e., thickness plateaus first, followed by volume, and finally surface area) the developmental stages at which they reach their peak differ between humans and macaques. In macaques, cortical thickness, volume, and surface area mature during infancy. In humans, however, there is a substantial delay: cortical thickness peaks during toddlerhood, volume during middle childhood around age 6, and cortical areas the latest during late childhood to adolescence at age 12. This indicates a prolonged gray matter development in humans compared to macaques. For both species, subcortical gray matter peaks in size at a similar mid-puberty, roughly at 4 years for macaques and 12 years for humans. Regarding white matter, humans peak during young adulthood, but no associated peak is found in macaques. Notably, ventricular data was available prenatally for macaques, but not for humans; accordingly, estimates from the human growth chart publication are annotated as a dashed line before birth in humans in Figure 3. Ventricular volume in macaques decreases dramatically before birth. After birth, it goes on to show a slow, progressive increase throughout the lifespan. In humans, a significant exponential increase is observed in the elder stage after 50 years. It’s important to note that the elder macaque samples in our study are limited (n=17 for age > 20 years, with the oldest being 33 years). While the typical lifespan of macaques in the wild is around 18-25 years^31^, those in captivity are reported to live up to 40 years. Therefore, the current sample might not fully capture the elder stage of brain atrophy or significant ventricular enlargement in the macaque lifespan.

### Timing and peak age of brain development across species

To characterize the critical brain developmental age, we calculated the regional peak age of brain growth for volume, surface area, and cortical thickness in macaques (Fig 4a, top). We also estimated peak age in humans using the previously established human charts^4^. In both humans and macaques, gray matter grows and reaches its maximal thickness first, then starts to thin, while the cortical surface continues to grow, with some regions expanding even after the volume peaks. Across the cortex, the peak age of volume, area, and thickness share similar spatial variability (human: r=0.27-0.64, p<0.001; macaque: r=0.40-0.44, p<0.001) except for the peak age of area versus thickness in macaque (r=0.03, p=0.80). In macaques, the visual cortex plateaus first, during infancy (< 3.5 months), while the motor cortex, inferotemporal sulcus, and inferior frontal pole plateau the latest. In humans, the thickness of the insular and cingulate cortex plateaus last during early childhood at 3.93 years of age, while expansion and thinning continue until adulthood at 26.6 years. To further compare the relative peak age across the cortex between humans and macaques, we utilized a previously developed cross-species transformation and mapped macaque peak ages to the human surface^33^. The differences in peak age rank between humans and macaques were calculated by subtracting the transformed macaque ranked map from the ranked peak age map in humans. Figure 4a (bottom) presents the differences (human versus macaque) of the ranks in volume, area, and thickness. Overall, across the cortex, insular and cingulate cortex matured relatively later than other regions in humans compared to macaques. This prolonged maturation period may provide an extended window of opportunity for these regions to support the development of more complex cognitive abilities unique to humans.

**Fig 4.**
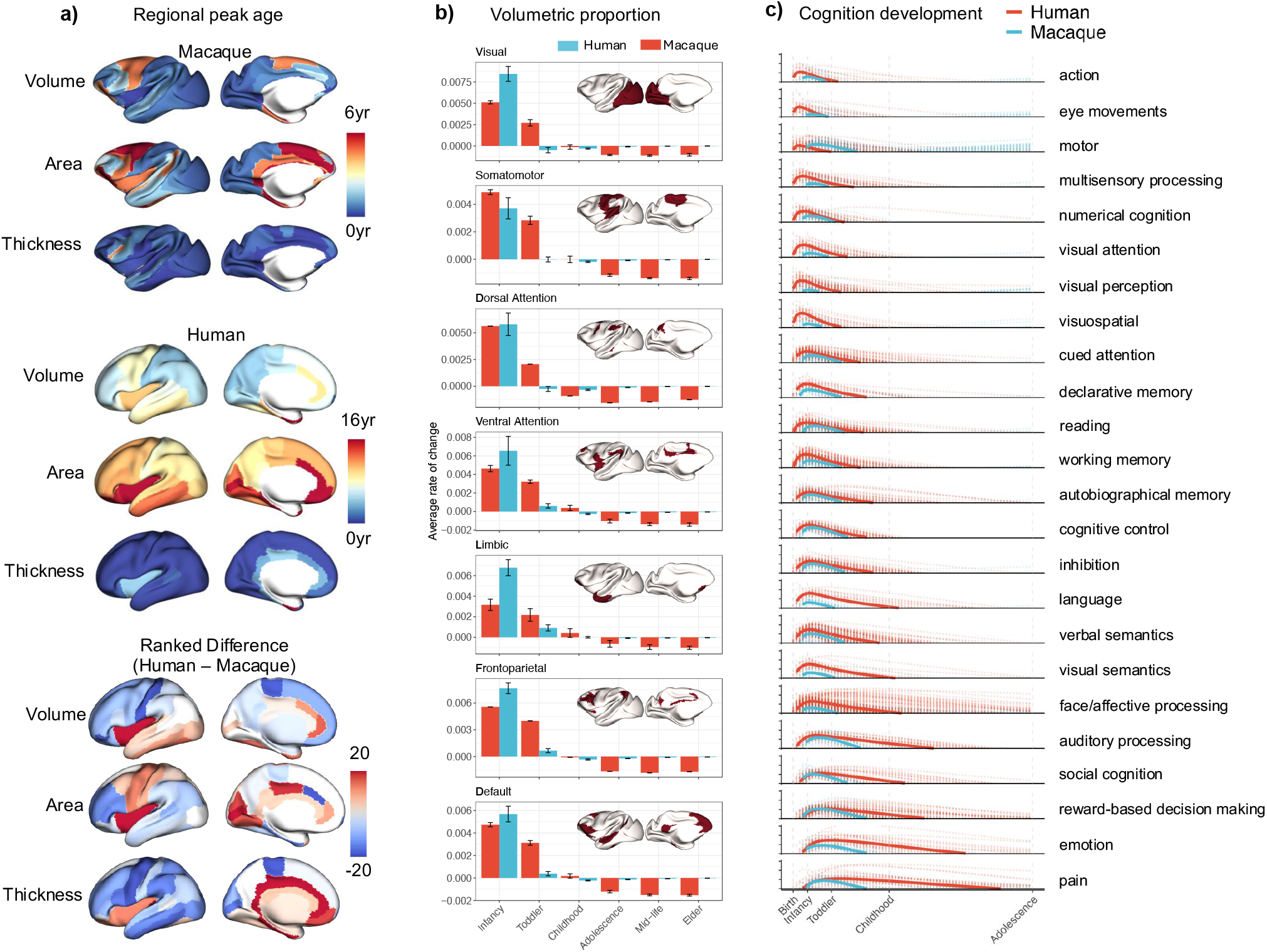
Regional peak age of morphological phenotypes, proportion of gray matter volume for each networks and cognition decoding of gray matter growth. **a**) Regional peak age of gray matter volume, surface area, and cortical thickness for human and macaque. Top: Macaque peak age maps. Middle: Human peak age maps from the human charts project^4^. Bottom: Relative differences in peak age rank between humans and macaques calculated by subtracting macaque peak age rank from the human peak age rank. Macaque peak age maps were ranked and transformed into human space for direct comparison. Red color indicates regions where humans mature later than macaques, compared to other cortical regional development; blue color indicates that macaque matures later in cortical development. **b**) Averaged proportional growth within networks at each developmental stage for both human and macaque. The developmental stages were approximately matched between human and macaque by averaging the respective proportions of total lifespan at each stage as noted in Fig 3). **c**) Decoded cognition development trajectories from birth to adolescence for human (0-18 years) and macaque (0-6 years). Regional growth maps of gray matter volume with a progressively increased interval (human: per 3 months; macaque: per month) were decoded using Neurosynth^32^ meta-analysis. Shaded lines represent individual terms associated with a topic, and solid lines represent the average.

At the network level (Fig 4b), early growth in human brain development is proportionally greater, while late growth reduction is also more pronounced. In contrast, macaques mature across all networks prenatally and exhibit relatively less change postnatally, with visual and motor networks showing a minimal decline from maturity to old age. In both species, the volume of higher-order networks decreases more than that of primary networks during the later aging process. This is much more notable in humans, whereas in macaques, the aging differences between higher-order and primary networks are not as apparent.

### Cognition development across species

To understand how differences in brain development relate to the formation of species-specific cognition and behavior, we conducted a Neurosynth^32^ meta-analysis utilizing volumetric cortical growth maps spanning birth to adolescence (Fig. 4c). Regional growth was calculated monthly for macaques and every three months for humans, with each region’s volumetric change computed as a proportion relative to the previous time point. Next, we transformed the macaque growth maps to human space using a previously established macaque-to-human transformation. Then, we decoded each growth map with a curated set of feature terms from Neurosynth, which encompass a spectrum of cognitive functions, including perception, action, and more abstract functions such as emotion and social cognition^34^. Finally, we summarized terms into 24 topics previously reported by calculating the averaged decoding score across terms for each topic at each time point (see Supplementary Information 5).

Sorting the decoded topics (i.e. the rows in Fig 4c) based on the age at which their associated cognitive function peaks in humans reveals a hierarchical sensory-association axis, a pattern that has been observed previously^34^. We observe short development windows from birth to infancy in regions associated with domain-specific sensory functions (e.g. “action”, “eye-movements”, “motor”, and “multisensory processing”), with both species approximately aligning in structural maturity trajectories relative to their natural lifespan. In humans, specifically, cognitive development is extended further on the age axis as we shift focus to regions implicated in complex representations of information and social cognitions (e.g. “face/affective processing”, “emotion”, “pain”, “social cognition”). It is worth noting that “pain” is widely recognized as a unique sensory and affective experience, characterized by an anatomically diffused phenomenon that is implicated in extensive areas of the limbic brain and prefrontal cortex^35^. In contrast, in macaques, Neurosynth scores between structural growth and high-order cognitive functions decline to zero much earlier in the lifespan. These findings concur with prior studies, suggesting that macaque and human brains are most structurally and developmentally different in higher-order brain regions^36^. This could represent the consequence of selective cortical enlargement and reorganization^37,38^ that evolutionarily separated us and macaques from our common ancestor.

### Brain Charts and Centile Scores Across Species

A crucial component of this work, and translational neurodevelopment as a whole, lies in directly comparing the brain and its development between humans and other primates. To advance the discovery and application of translational research, we developed an interactive open resource (https://interspeciesmap.childmind.org) for the cross-species comparison of global and regional brain MRI phenotypes over the human and macaque lifespans following alignment based on developmental milestones (Fig. 5). This online platform integrates previously established human brain charts and corresponding centile scores with results presented here. It also utilizes prior macaque-human cortical alignment to display homologous brain regions between species. The normative model with centile trajectories, stratified by sex and annotated with peak growth velocities and their maximums are available for quick and interactive explorations for both humans and macaques. It aids NHP researchers in benchmarking the brain structure with limited animal samples for control groups and identifying the matched MRI-driven milestones to better understand the biological processes of brain development.

**Fig 5.**
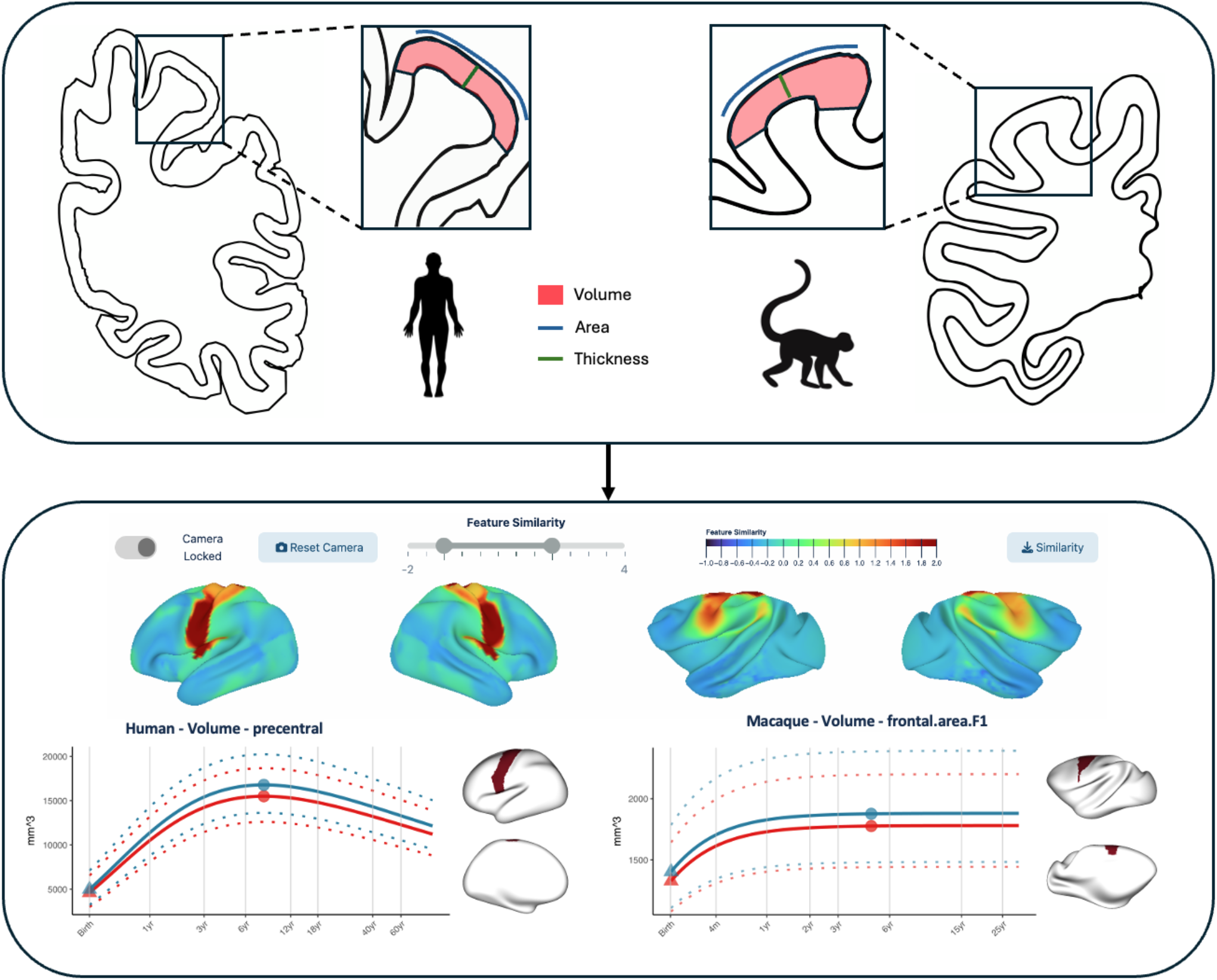
Demonstration of interactive cross-species brain charts. Top panel depicts normative models of the morphological measurements derived from MRI for macaque and human. Bottom panel depicts homologous regions between human and macaque on cortical surface maps with associated growth trajectories. Growth trajectories for each region are shown on a log scale, including centile trajectories, stratification by sex, and annotations for its maximum (circles) and peak growth velocity (triangles).

## Discussion

The present study established lifespan growth charts for brain structure in macaques that cover late gestation, infancy, juvenility, adolescence, early adulthood, mid-adulthood, and late adulthood. Importantly, we compared macaque brain development trajectories with the previously established human brain charts to understand neurodevelopmental resemblance across species. Overall, both human and macaque brain development follows an inverted-U-shaped trajectory; however, each species exhibits a specific developmental timeline. Unlike humans, the peak growth rate of gray and white matter in macaques occurs prenatally, resulting in a more mature total brain volume at birth, with 50% to 75% lifetime size. In contrast, humans are born with only 25-30% of their adult brain size, and experience a prolonged growth period extending into adolescence, allowing more time for experience to shape higher-order cognition and social learning. Additionally, we provided sex-stratified, centiled normative brain trajectories through an interactive open-resource platform, supporting cross-species comparison of brain development between humans and macaques. With the rise of open-data sharing initiatives in the NHP community, these brain charts can be further refined to facilitate benchmarking of NHP brain development, potentially allowing for more flexibility in sample size requirements for future NHP research.

Based on the largest macaque brain lifespan sample to date (1,522 scans from over 1,024 macaques aggregated from 23 sites), the resulting lifespan trajectories align well with existing literature (sample size ranging from 7-66 macaques) and characterize developmental milestones of the cerebrum in macaques^39,19,40,41,18,42^. Total GMV peaks at 0.74 years (∼9 months, CI: 0.65 - 0.86 years), coinciding with previous studies that have also reported that GMV reaches its maximum size around 1 year of age, with no significant maturational change after the first 1-2 years^39,43,18^. Prior to GM volume plateau, cortical thickness matures earlier, reaching its peak around 3 months of age–during infancy–then gradually declining to adult thickness, with the trajectory itself resembling an inverted-U shape. This result is remarkably consistent with other longitudinal neuroimaging studies^44^. It also aligns with the timing of synaptogenesis, where synaptic density increases significantly prenatally to 3 months of life, followed by synaptic pruning that occurs into adulthood^26–28^. Similar to the GMV charts, the peak growth rate of WM volume in macaques also occurs during gestation, reaching over 50% of adult size at birth in macaques. This may reflect the rapid increase in myelinated axons prenatally compared to humans^19,40,41^. After birth, the increase in white matter volume to adult size is substantially longer than the increase in gray matter and total brain volume in both species^19,45^. After infancy, growth velocity in humans is greater than in macaques, progressively growing to a greater percentage with increasing age.

Regarding subcortical gray matter, previous studies reported sGMV in macaques gradually increasing up to 3-4 years of age in early developmental cohorts^18,42^ and decreases in older cohorts^5^. Our study narrowed these estimates, suggesting that peak maturation occurs around 4.39 years (CI: 3.70yrs - 5.75yrs), occurring post GMV peak, during adolescence. Interestingly, in humans, total subcortical GM volume also peaks later than the GMV short after sexual maturation, followed by a gradual decline^4^. Given that subcortical structures have been highly conserved throughout primate evolution^46,47^, it is not surprising that both species share similar growth phases. However, aging processes for specific subcortical nuclei differ between humans and macaques. While both species experience volume declines in caudate, putamen, and globus pallidus^48,49^, the hippocampus shows notable differences^49^. In humans, hippocampal volume decline is a hallmark of aging, with atrophy accelerating around middle age^50^. In contrast, such a reduction in the size of the hippocampus has not been observed in aging macaques and does not appear to be related to cognitive memory impairment in older macaques^48,51–53^. Similarly, in chimpanzees, hippocampal volume does not show significant age-related changes either^54^. This suggests that hippocampal changes seen in humans may be unique in aging when compared to other primates^48,51–53^. Additionally, a notable difference in aging trajectories is also observed for total ventricular volume between species. Unlike humans, who have significant exponential ventricular enlargement after 50 years, macaques show a steady increase throughout their lifespan. Although our current sample (up to 33 years) might not fully represent the elder stage of brain atrophy in the macaque lifespan—since macaques in captivity can live up to 30-40 years—our findings suggest species-specific aging mechanisms. It’s also important to consider that the relative lifespan of macaques in captivity may be more closely aligned with the concept of “healthspan” in humans, with comparable humane endpoints being determined as the point when the animals can no longer perform daily activities independently^3,55–57^. These divergences potentially reflect a greater vulnerability of humans to age-related neurodegeneration diseases such as Alzheimer’s and Parkinson’s^48^, underscoring the importance of the large-scale macaque charts in understanding human brain development and aging.

In general, therefore, macaques are born more mature and develop at a rate 3-4 faster than humans^18,30^. In contrast, humans exhibit a prolonged growth period extending into adulthood, indicating neotenous neurodevelopment within primate lineage. In macaques, the growth velocities for volume, thickness, and area peak before birth, while in humans, only cortical thickness shows significant growth prenatally. After birth, the human neocortex grows at a slower rate than that of macaques, retaining plasticity for an extended period of time. Importantly, the rate of brain maturation varies across regions. The most pronounced delays in human brain maturation occur in the anterior cingulate cortex (ACC) and insula (Fig 4), both regions highly involved in cognitive, emotional, affective and pain processing in humans^58–60^. The ACC is a hub that integrates neural signals from the limbic system, hippocampus, and a wide range of cortical areas, supporting many cognitive and emotional functions such as self-regulation, social interactions, reward, and motivation^59,61–64^. The insula is also a multifunctional area, supporting functions such as risky decision-making, emotional subjective feelings, and social interaction^65,66^. Recent fMRI studies have also revealed links between the anterior agranular insula and self-reflection^67^, interoceptive attention^68^, and awareness of bodily and emotional states^69^, highlighting its crucial role in human consciousness^65^. While the insular region in macaques also contains similar dorsal cognitive and ventral emotional subdivisions, macaques generally lack self-awareness and do not typically recognize themselves in mirrors^70,71^, though some may pass mirror tests with extensive training^72,73^. Prior literature has also shown that the anterior agranular insula is among the most significant expansions in humans compared to chimpanzees^74^. Although the current parcellations are too coarse for modeling age-related changes in subregions of the ACC and insula, the striking difference in the maturation timeline between species supports their unique roles in human behavioral evolution.

Notably, other high-order association regions, such as frontoparietal and default mode networks, also exhibit neoteny, with protracted maturation in humans compared to macaques. However, unlike the insula and ACC, these regions are not the last to mature anatomically. After their gray matter volume reaches adult size, functional connections continue shaping brain modularity, increasing structure-function coupling, and transitioning from a sensory-centered schema to a hierarchically distributed organization^29,75–77^. To characterize the critical development stage and its associated behavior, we further decoded the incremental growth maps with a NeuroSynth meta-analysis. For both species, the most developmental similarities in the incremental growth maps lie in basic primary sensory related behavior tasks, such as those associated with “action” and “visual perception”. However, as topics begin to represent regions associated with high-order functioning (“decision-making”, “emotion”, “affective processing”, “pain”), humans exhibit much longer growth timelines. This reinforces the hypothesis of a protracted maturational trajectory in humans, disproportionately favoring the transmodal/association end of the hierarchical sensory-association axis^34^.

Interestingly, not all regions display prolonged development timelines in humans. The precentral motor cortex in macaques grows more slowly compared to humans, with its volume and area fully reaching adult size between 4-9 years of age, during late adolescence, and extending to adulthood. The delayed maturation of the precentral cortex provides increased plasticity in macaque, allowing the refinement of learning and adaptation of fine motor skills^78^. Our meta-analysis echoed this finding, showing that the “motor” function had an extended growth timeline in macaques compared to humans. Taken together, the respective delayed maturation of specific regions in each species reflects species-specific adaptation to the environmental experience throughout evolution.

Leveraging open science and data sharing, this study has collectively aggregated the largest macaque MRI dataset to date, spanning an age range that covers the lifespan. The resulting global and regional charts provide the first normative model, setting a benchmark for future macaque brain development research. This resource could potentially reduce the sample size requirements for studies investigating brain plasticity, vulnerability to neuropsychiatric disorders, and aging-related diseases, thereby supporting the ‘3R’ (Replacement, Reduction, and Refinement) principles of ethical animal research. It’s worth noting, however, that most of the samples in this study are cross-sectional, which might not capture subtle developmental nuances attainable from multiple within-subject scans. For example, longitudinal studies have revealed a unique period of no net gray or white matter growth between 6-9 months. This period, reported in several studies, could be related to weaning, stress of newfound independence, or increased physical activity during this period. Probing such subtle intricacies warrants further validation through additional biometric and behavioral measurements. In late adulthood, our fitted trajectory showed no decline in total WM volume, contrasting with previous reports of age-related WM reductions in older macaque^5,6,49^. This discrepancy is likely due to the limited sample size for macaques older than 20 years (N=17, with only one male among them). As more open data-sharing initiatives emerge in NHP research, we anticipate the inclusion of more high-quality MRI data in the future, maximizing the reuse of NHP resources and enabling cross-species comparative models for studying the human brain. The GAMLSS modeling framework lends itself to the continued iterative aggregation of such data, allowing for the non-linear, age-related modeling of higher-order parameters (variance and skewness). To facilitate further investigations using brain chart resources in humans and macaques, we have provided cross-species whole-brain cortical mapping and lifespan normalized centile scores for humans and macaques, available at https://interspeciesmap.childmind.org/.

## Contributions

T.X. and M.P.M. conceptualized and designed the study. S.A. and T.X. conducted the analyses, drafted and wrote the manuscript. S.A. and J.R. organized and preprocessed the data. R.V.W. developed and implemented the website. Z.W., N.B.E., and A.F. assisted with data organization. T.X. supervised the overall project. R.A.I.B. and J.S. provided support with statistical analyses. All other authors were involved in one or more of the following tasks: data acquisition and collection, organizing datasets, analyzing results, interpreting findings, editing the manuscript, or reviewing the manuscript.

## Acknowledgments

T.X. was supported by the National Institutes of Health (NIH) RF1MH128696 and P50MH109429. M.P.M was supported by NIH R01MH111439, P50MH109429, MH124045, MH230482, and MH109429. R.A.I.B. was supported by the Academy of Medical Sciences, HDR UK. J.S. was supported by NIH R01MH134896, R01MH132934, and R01MH133843. A.F.A.-B. was supported by NIH R01MH134896, R01MH132934, and R01MH133843. Data were curated and analyzed using the Pittsburgh Supercomputing Center (BIO220056) through the Advanced Cyberinfrastructure Coordination Ecosystem: Services and Support (https://access-ci.org). We acknowledge the contribution from open datasets used in this study: the PRIMatE Data Exchange (PRIME-DE) (https://fcon_1000.projects.nitrc.org/indi/indiPRIME.html) and the UNC-Wisconsin Neurodevelopment Rhesus Macaque MRI Database (https://www.nitrc.org/projects/uncuw_macdevmri). We thank Dr. Leslie Ungerleider and Paula L. Croxson for their contributions by providing the NHP data. We also express our deep appreciation to all the facility staff who assisted with the NHP data collection and acquisition.

## Methods

### Dataset and Preprocessing

The nonhuman primate (NHP) data was aggregated from the PRIMatE-Data Exchange (PRIME-DE) consortium, UNC-Wisconsin Neurodevelopment Rhesus MRI Database^1^, and additional development studies. Refer to Supplementary Information 1.1-2.1 for details on individual study protocols, scanner manufacturers, and demographics. We preprocessed T1w and available T2w MRI images using ‘deepbet’ and ‘nhp-abcd-bids-preprocessing’^2–4^ pipelines. Briefly, the data was first denoised using ANTs to remove the ‘salt-and-pepper’ noise, followed by skull stripping using ‘deepbet’, segmentation, surface reconstruction in FreeSurfer^5^, and registration to the MacaqueYerkes19^6^ template. Additional segmentation templates from the early development stage were added into the ‘nhp-abcd-bids-preprocessing’ pipeline to successfully segment infant macaques with age < 0.33 years. Quality assessment figures and metrics were generated for each preprocessing step for visual inspection and quantitative quality control (see Supplementary Information 1). In total, 1,024 rhesus macaques (female=470, male=554, age range -0.227 - 30.64 years) with 1,522 unique scans from 23 research sites were included in the analyses. We then extracted global and regional volume, area, and thickness measurements utilizing the Markov^7^ parcellation to define regions across the cortices. ComBat was applied to global and regional measurements to adjust for potential batch effects that might arise from variations in the segmentation stage (see Supplementary Information 1.5).

For cross-species comparisons, we utilized the previously established normative charts from the human study^8^. The GAMLSS modeled trajectories for global measurements (i.e. total volume, total area, and mean thickness) as well as the regional metrics defined by the Desikan-Killiany^9,10^ parcellation is available at https://github.com/brainchart/Lifespan.

### GAMLSS and Growth Charts

To model age-related changes in brain structure across the lifespan with an aggregate dataset, we employed the Generalized Additive Model for Location, Scale, and Shape (GAMLSS)^11^. This approach allows us to fit non-linear growth curves with a wide range of distribution families that incorporate the variations in mean, variance, skewness, and kurtosis. A recent large-scale study has established human brain growth charts from over 100,000 scans. Following this approach, we utilized GAMLSS to model nonlinear sex-stratified developmental trajectories on the macaque lifespan data. The generalized gamma distribution family, fractional polynomials, has been shown to be the most suitable outcome distribution for modeling brain growth curves across the lifespan, as it effectively captures the non-linear changes, heteroscedasticity, and skewness present in the data^8^. Therefore, fractional polynomials were optimally selected to model the growth trajectories with non-linearity in *μ* for each of the global and regional MRI measurements. Specifically, the following GAMLSS model is specified:

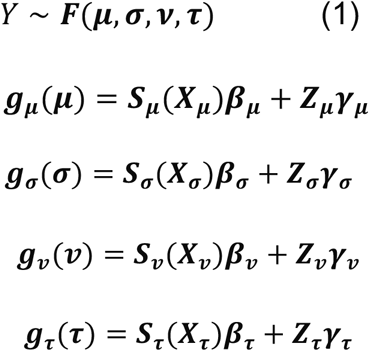

The dependent variable, Y is modeled by a probability function *F*, commonly parameterized by distribution parameters: mean (μ), variance (**σ**), skewness (**ν**), and kurtosis (**τ**). The location and scale of the distribution are characterized by mean and variance while the shape is characterized by skewness and kurtosis. Each parameter can be modeled as a linear function of explanatory variables through a link function ***g***(). ***X*** and ***Z*** are design matrices for fixed effect ***β*** and random effects ***γ*** respectively. Non-parametric smoothing functions applied to independent variables for each of the four parameters are denoted as *S*. Similar to the prior work in humans, we used fractional polynomials as smoothing functions for the age effect in ***X*** and estimated the associated coefficients ***β***. For each dependent variable, we fit all possible fractional polynomial models iteratively from a standard set of powers, *p* ∈ {−2, −1, −0.5,0,0.5,1,2,3}, searching for the most appropriate subset of powers that optimize the model fit. The model with the lowest Bayesian information criterion (BIC) was selected as the final model for each global and regional brain metric. Of note, while the sample size is considered a large dataset in NHP studies, the age coverage from multiple datasets might be inadequate to model the non-linearity in variance (*σ*). As such, the searching algorithm was only estimated for the first order of distribution parameters (μ).

In line with the previous study^8^, ‘site’ was also included in the model as a random effect, estimating mean (***μ***) to account for batch effort from the study site. Specifically, beyond the fixed intercept, observations that varied across ‘site’ were modeled by the random effect of the intercept, assuming the site variation follows a normal distribution *γ∼N*(0, *δ*^2^). Therefore, the potential batch effect - site-specific deviation - was taken into account in GAMLSS and estimated by the random effect term in the model. See further validation analyses in Supplementary Information 3.1-3.3. We also estimated the study-specific model for each site by removing the ‘site’ effect from the model (Fig 1, Fig SX), which characterized the brain growth curves for each site.

Finally, we identified the optimal fractional polynomial models for each of the global and regional metrics. The models for the global metrics are as follows:

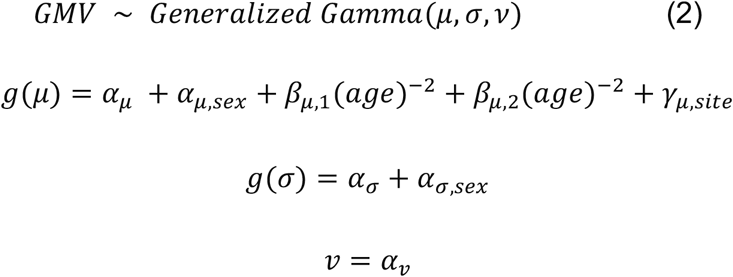

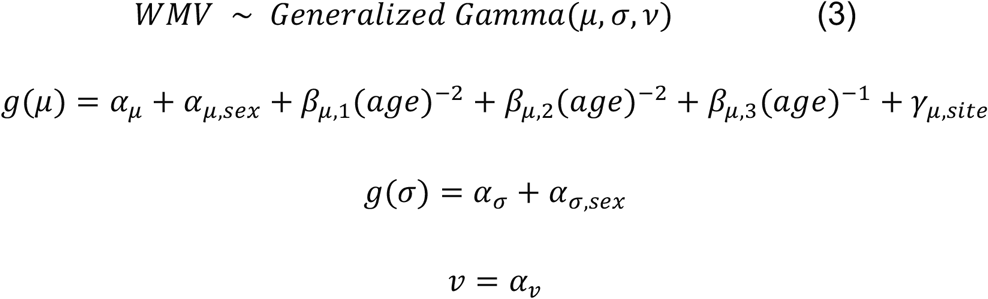

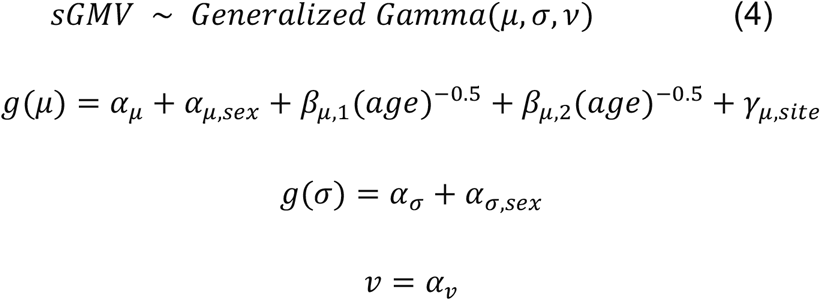

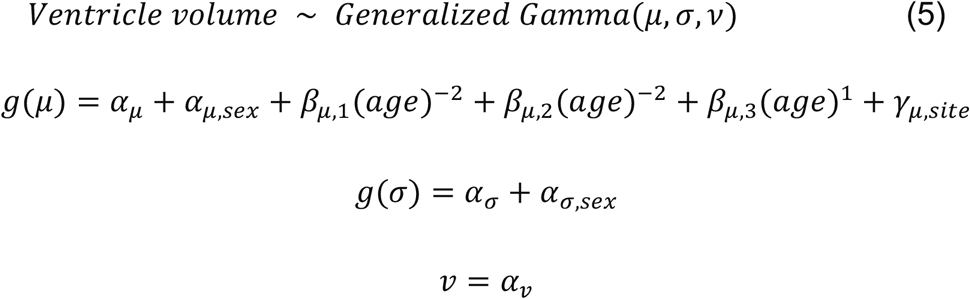

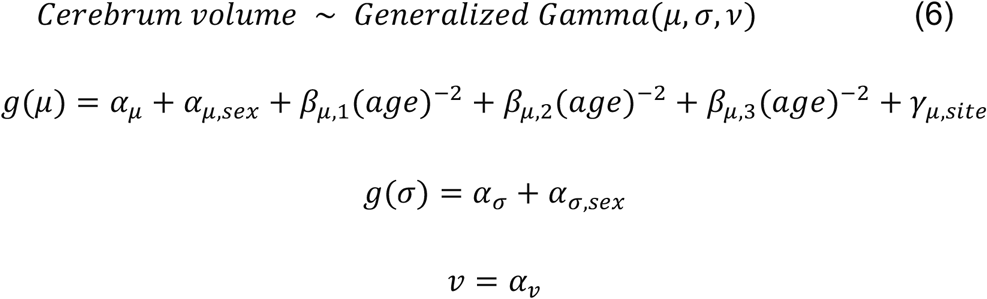

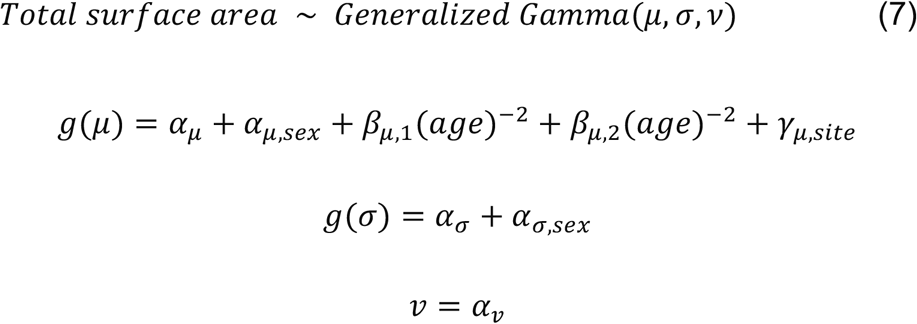

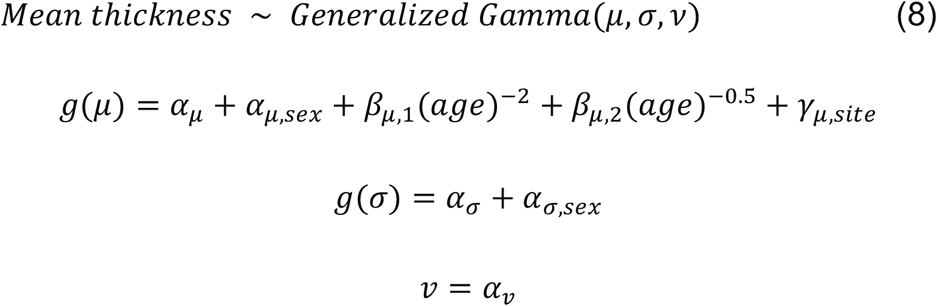

The term *α*_μ_ corresponds to the fixed effect of the intercept, which is modeled for mean (*μ*), variance (***σ***), and skewness (***ν***); The fixed effect of sex, *α*_*μ*,sex_ is modeled for the mean (*μ*), variance (***σ***); the term *γ*_μ_,_site_, representing the random effect of study ‘site’, is also included in modeling mean (μ).

To validate our models and calculate confidence intervals of the resulting trajectories and peaks of growth rate and age, we performed a bootstrapping analysis, consisting of 1,000 refits with replacement. Trajectories for global and most regional metrics are robust to this analysis, with several instabilities found in smaller regional estimates. Details for this analysis and the results can be found in Supplementary Information 3.

### Cross-species transformation

To compare spatial patterns across species, we utilized previously established functional alignment to transform resulting maps between macaques and humans^12^. This method provides a vertex-to-vertex surface deformation across species, constructed based on the matched low-dimensional representations of functional connectivity in humans and macaques. Code and transformation are available: https://github.com/TingsterX/alignment_macaque-human.

### Determining developmental milestones across species

To quantify and compare developmental milestones, we used GAMLSS trajectories to estimate ages at which regional and global measurements peak in their size and growth rate. Growth rates were determined by taking the first derivative of the median trajectory (50th centile), with the peak growth velocity identified at its maximum. Similarly, the age at which a tissue type or region peaked in volume, area, or thickness was determined by the point at which the median trajectory reached its maximum. In some cases, particularly in smaller regions, GAMLSS trajectories were observed to plateau in growth until the end of the lifespan. In these instances, the earliest point of the plateau was recognized as the peak age (See Supplementary Information 4 for details).

To directly compare the order of regional maturation between species, peak ages were first projected back to the cortical surface. We then transformed the macaque maps to human surface space using the cross-species alignment described above. The macaque-to-human peak age maps were re-parcellated using the Desikan-Killiany^9^ atlas to match the regional human peak age maps. Following, we ranked the peak age from earliest to latest maturation time within species and directly subtracted the resulting ranked maps (human - macaque) to generate a cross-species difference map, representing the contrast in relative maturity of regions across the cortices.

### Neurosynth Meta-analysis decoding of growth maps

To identify critical developmental stages and associated cognitive function, we conducted a meta-analysis using the Neurosynth^13^ database. We first calculated incremental growth maps, dividing the developmental period from birth to early adulthood (macaque: 6 years, humans: 18 years) into time windows of 3-month and 1-month intervals in humans and macaques, respectively. For each interval, we calculated the volumetric change of cortical gray matter regionally, resulting in 72 growth change maps in each species. Following, we decoded each of the human growth maps using the Neurosynth meta-analysis database with the Brainstat^14^ software package, computing the spatial correlations (i.e. r-score) between our growth maps and the Neurosynth term activation maps. As activation maps in the database are all in human reference space, we projected macaque growth maps onto the human surface before decoding the macaque-to-human map accordingly. Finally, we summarized feature terms into 24 topics previously reported^15^ (e.g. feature terms such as ‘sight’, ‘vision’, and ‘eye’ are grouped into the topic ‘eye movements’). The r-scores of terms from the same topic were averaged for each time window within species. In Fig 4, the dotted curve represents the connected scatter plot of the r-score for each feature term, and the solid curve represents the averaged score across terms for each topic. Each curve indicates the cortical developmental window for a specific cognitive function. Additionally, we also used GAM models to fit the r-scores across terms to generate the topic curve (Supplementary Information 5.1).

